# Dissecting the Contribution of Transposable Elements to Interphase Chromosome Structure

**DOI:** 10.1101/2024.10.25.620372

**Authors:** Liyang Shi, Zhen Xiao, Isaac A. Babarinde, Fu Xiuling, Gang Ma, Zhouqi Huang, Li Sun, Xuemeng Zhou, Jiangping He, Alexander Strunnikov, Andrew P. Hutchins

**Affiliations:** Department of Systems Biology, School of Life Sciences, Southern University of Science and Technology, Shenzhen, Guangdong, China; Laboratory of Inflammation and Vaccines, Shenzhen Institute of Advanced Technology, Chinese Academy of Sciences, Shenzhen, Guangdong, China; Guangzhou National Laboratory, Guangzhou, Guangdong, China; Guangzhou Institutes of Biomedicine and Health, Guangzhou, Guangdong, China

**Keywords:** Transposable elements, 3D genome, transcription factors, epigenetics, SMARCA4, BAF complex

## Abstract

Transposable elements (TEs) occupy nearly half of the human genome and play diverse biological roles. Despite their abundance, the extent to which TEs contribute to three-dimensional (3D) genome structure remains unclear.

To investigate this, we developed te_hic, a computational pipeline that integrates genomic data with chromatin interactions. Our analysis reveals that TE sequences are responsible for 3D genome structure in interphase nuclei. This phenomenon is mediated by the recruitment of specific transcription factors to TEs, which enables both the formation and disruption of chromatin contacts. We computationally identified known contact-forming proteins (CTCF, RAD21, SMC3) and breaking proteins (RNF2), as well as novel candidate contact-formers (SMARCA4, MAFK). Knockdown of the predicted contact-formers *SMARCA4* and *MAFK* decreased contacts and loops at and between TEs. Notably, *SMARCA4* knockdown selectively reduced short-range contacts, highlighting its role in maintaining 3D genome structure mediated by TE binding.

Overall, our findings demonstrate that TEs are crucial determinants of 3D genome organization in mammalian cells.

**Key findings:**

- TEs determine 48% of the 3D genome structure alone, and 70% of heterotypic contacts.
- A/B compartments, TADs, and loop signatures can be retrieved using TE-mapped reads only.
- TFs can be divided into contact formers and breakers at TEs.
- SMARCA4 and MAFK build chromatin contacts between TE sequences.

## Background

DNA is tightly packaged in the human nucleus and is wrapped in complex three-dimensional structures that are involved in regulatory processes [1, 2]. DNA and chromatin are arranged in hierarchical levels [3]. In roughly descending order of size: A/B compartments, that roughly correspond with euchromatin and heterochromatin, megabase-sized domains called topologically-associated domains (TADs) which can be further subdivided into smaller sub-TADs [1], chromatin loops, which are driven by CTCF and Cohesin [3], and direct contacts between specific loci, such as enhancer-promoter contacts [4]. These structures have been identified in multiple species, although they are present at greater or lesser degrees [1]. These chromatin structures have also been implicated in a range of biological processes, including developmental processes and human disease [5, 6]. However, the principles that govern the formation of these domains remain only partially explained and no comprehensive model describes the 3D genome.

Nearly 50% of the human genome consists of fossilized transposable element (TE) sequences [7]. TEs are genetic elements that are capable of duplicating themselves across the genome, although the vast majority of TEs are inactive [8]. TEs are involved in a wide range of cellular and biological processes [8], including being transcribed as parts of non-coding transcripts [9], as RNA-binding protein binding sites [10, 11], in early embryonic development [12, 13], and human disease [14].

A role for TEs in 3D genome structure has been suggested by several studies [15, 16]. One clear example is the role of the key loop-forming protein CTCF. Several types of TE contain a CTCF binding site and have driven the evolutionary redistribution of CTCF binding sites to shape 3D genome structure [17–19]. The best-characterized example is the expansion of the rodent-specific SINE B2 TE that contains a CTCF binding site which has extensively duplicated in rodents to introduce novel CTCF binding sites that have altered the 3D genome structure [18, 20, 21].

Other classes of TEs have been implicated in a more general way. Human pluripotent stem cells (hPSCs) express RNAs from endogenous retroviruses (ERVs) [9], and several HERV-Hs sit at the border of TADs and genetically deleting them removes TAD boundaries and merges nearby TADs [22]. MIR family TEs can form insulator boundaries between TADs [23], and form enhancer-like elements that bridge the TE sequence to promoters [24]. LINE-2, Charlie, and MaLR TEs are associated with mammalian TAD boundaries [25]. Systematic analysis has also indicated LINE and SINE family TEs demarcate genome compartments and chromatin states [21, 24, 26]. These studies illustrate how TEs are involved in 3D genome formation. However, except for CTCF, mechanisms that form TE-mediated 3D contacts are lacking [27]. As TEs are rich in transcription factor (TF) binding sites [12, 28–30], it seems likely that TFs bound to TEs are responsible for forming the 3D genome structure. We propose that some classes of TFs are capable of breaking and forming DNA contacts at specific families of TEs.

Here, we used the comprehensive TF binding data available in human and mouse pluripotent stem cells (PSCs), coupled with Hi-C data to explore the role of TFs bound to TEs in 3D genome organization. Our results show that TEs dominate the organization of the 3D structure of the genome and are responsible for driving homotypic TE-to-TE contacts. We divided TFs into three main classes: those that utilize TEs to drive 3D genome formation, those that are neutral, and a third class that breaks 3D contacts at specific TEs. Based on these analyses we show that SMARCA4 and MAFK are involved in promoting chromatin contacts at TEs.

## Results

### Process Hi-C data to include transposable elements

Hi-C is a powerful method to elucidate the 3D structure of the genome [31]. However, in typical Hi-C experiments TEs are under-represented, as reads mapping to TEs are either actively removed or indirectly removed as low-quality reads (**Figure 1A**). To investigate TEs in 3D genome structure, we implemented a Hi-C processing pipeline that specifically considered TE-mapped reads (https://github.com/oaxiom/te_hic) (**Figure 1B**). The pipeline is similar to the approach taken in the HiC-Pro workflow [32]. Briefly, reads from Hi-C data are independently aligned to the genome using bowtie2 [33]. Paired reads are then filtered based on certain criteria: (1) Both pairs must be mapped to the genome; (2) Both pairs must have a mapping quality of at least 10; (3) Paired reads must be at least 2 kbp apart. Multi-mapped reads are assigned to the single best mapping locus. This contrasts with HiC-Pro, which has a default quality score of 0 but removes all multi-mapped reads. The ENCODE Hi-C pipeline uses a minimum mapping quality of 30, and also discards all multi-mapped reads [34]. te_hic then assigns each end of the paired reads to a genomic locus, either inside or outside a TE producing a BEDPE file that contains a list of all qualified paired reads and if the read is inside or outside of a TE sequence. Finally, te_hic generates four raw count matrices of chromosome contacts: (1) All qualified reads (2) Both reads do not map to a TE (no-TE to no-TE) (3) one paired-end read maps to a TE (TE to no-TE) or (4) both paired reads map to a TE (TE to TE). The resulting matrices are suitable for downstream analysis using typical Hi-C processing and normalization tools. In our work, we used ICE (iterative correction and eigenvector decomposition) [35] to balance the resulting matrices for downstream analysis.

**Figure 1.**
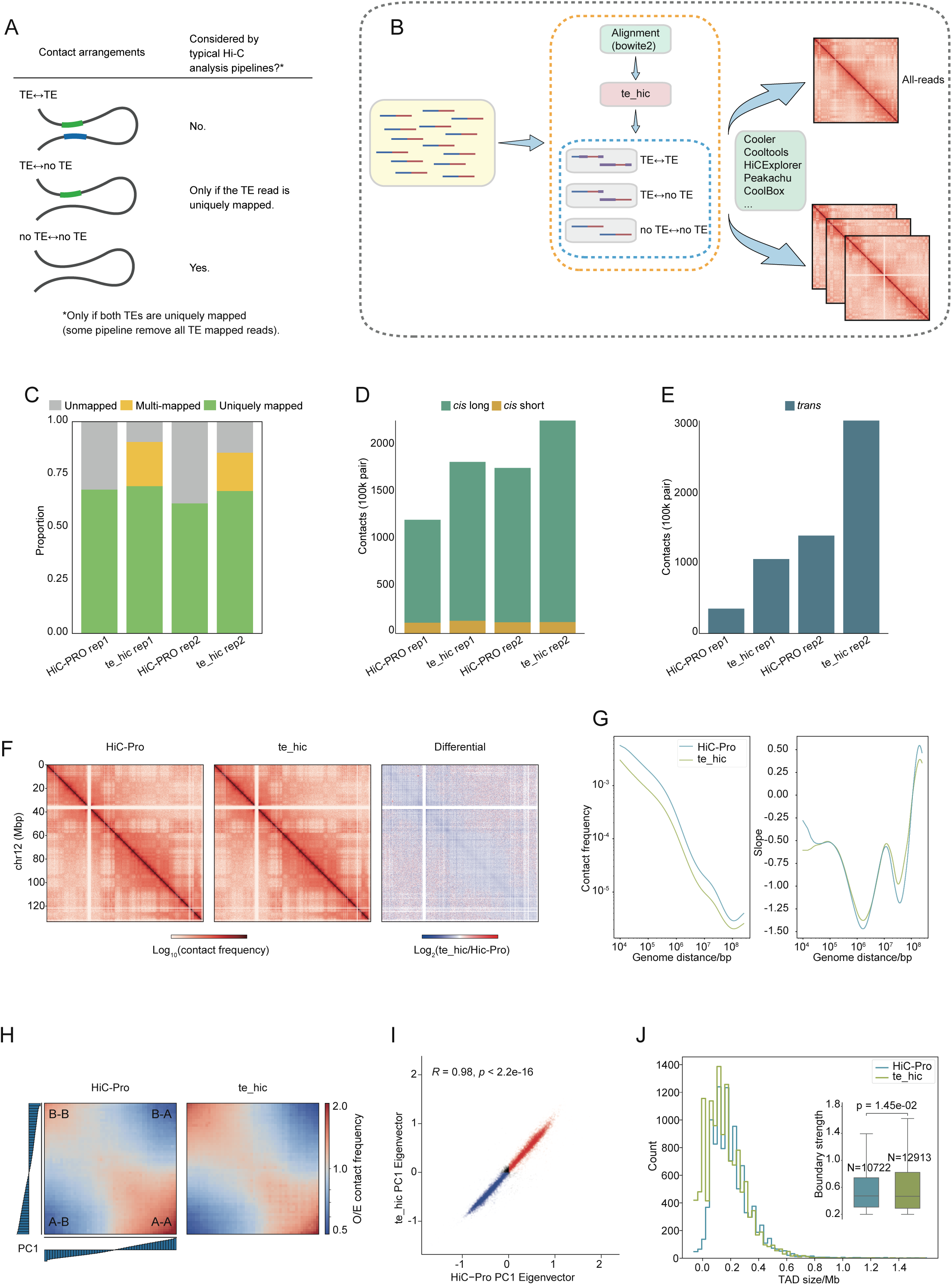
A Hi-C pipeline that integrates TEs. (A) Schematic of possible TE arrangements in 3D chromatin structure. (B) Schematic of the te_hic pipeline. The implementation is available at https://github.com/oaxiom/te_hic. (C) Bar plot showing the proportion of unique and multi-mapped reads from Hi-C data in GSE52457 [36] (D) Bar plot showing the breakdown of the detected paired-read contact types: *cis*-short range interactions (<20 kbp), and *cis*-long range interactions (>20 kbp), where both reads are on the same chromosome. (E) Barplot of *trans* chromatin contacts, where both reads are on different chromosomes. (F) Contact heatmaps and differential contact heatmap of chromosome 12 from the matrix generated by HiC-Pro versus the ‘all-reads’ matrix from te_hic at 150k resolution. Log2 comparison between two matrices was performed using ‘hicCompareMatrices’ of HiCExplorer [81] with default parameters. (G) Line plots showing contact frequency as a function of distance of HiC-Pro matrix and the te_hic-all matrix. (H) Saddle plots showing genome-wide average observed vs. expected (O/E) interaction frequency and A/B compartmentalization of HiC-Pro matrix and the te_hic-all matrix. The left and below bars indicate the genomic bins sorted by PC1. (I) Scatter plot showing the A/B compartment scores for each pixel in the HiC-Pro matrix versus the te_hic ‘all-reads’ matrix. (J) Histogram of TAD domain sizes in megabase pairs as determined using matrices from HiC-Pro and te_hic. Boxplot of strength of TAD boundaries, and number of boundaries is indicated (subplot).

We validated the overall approach of te_hic by comparing it with HiC-Pro [32]. To this end, we reanalyzed a deeply sequence hPSC Hi-C dataset [36] using both tools and compared the ‘all-reads’ matrix produced by te_hic to the matrix produced by HiC-Pro. Both tools utilized bowtie2 as their default aligner and te_hic retained more reads as compared to HiC-Pro (**Figure 1C**). We next measured overrepresented chromatin contacts relative to over-enrichment based on a Poisson distribution for the number of contacts between all pairs of genomic loci. Overall, although te_hic did not discover more *cis*-short (>5kbp <20kbp), te_hic tended to detect more *cis-*long (>20kbp) and *trans* contacts, for the same replicate samples (**Figure 1D, E**). A visual inspection of the all-reads matrix for chromosome 12 indicated little difference overall, although te_hic tended to report a decrease in signal (**Figure 1F**). This was reflected in a global decrease in contact frequency as a function of distance (**Figure 1G**). A/B compartments were nearly identical between HiC-Pro and te_hic matrices (R=0.98, p=2.2e-6; **Figure 1H, I**). Boundary strength was also broadly similar between HiC-Pro and te_hic (**Figure 1J**). Finally, TAD sizes were also similar between HiC-Pro and te_hic (**Figure 1J**). Overall, the data indicates that the incorporation of TE reads and multi-mapped reads into Hi-C data analysis does not disrupt the broad pattern of 3D features and does not lead to drastically different analytical outcomes. This will then allow us to dissect the contribution of TEs to 3D genome structure.

### Transposable elements underly 3D chromatin structures

We next explored the TE-specific matrices produced by te_hic. We first dissected the contribution of TEs by measuring the proportions of read pairs in the Hi-C data with both reads mapping to a TE (TE-to-TE), with only one end mapping to a TE (TE-to-no-TE) and both reads not in a TE (no-TE-to-no-TE). 70% of all the mapped reads had at least one read pair inside a TE and for only 22% of reads both ends mapped outside of a TE (**Figure 2A**). A Monte Carlo simulation using random genome coordinates indicated that this proportion is biased towards TE sequences, as the expected contact percentages by chance alone are: TE-to-TE = 30.4%; TE-to-no-TE = 46.7%; and no-TE-to-no-TE = 22.8%. This indicates TE sequences contribute disproportionately to 3D genome sequence.

**Figure 2.**
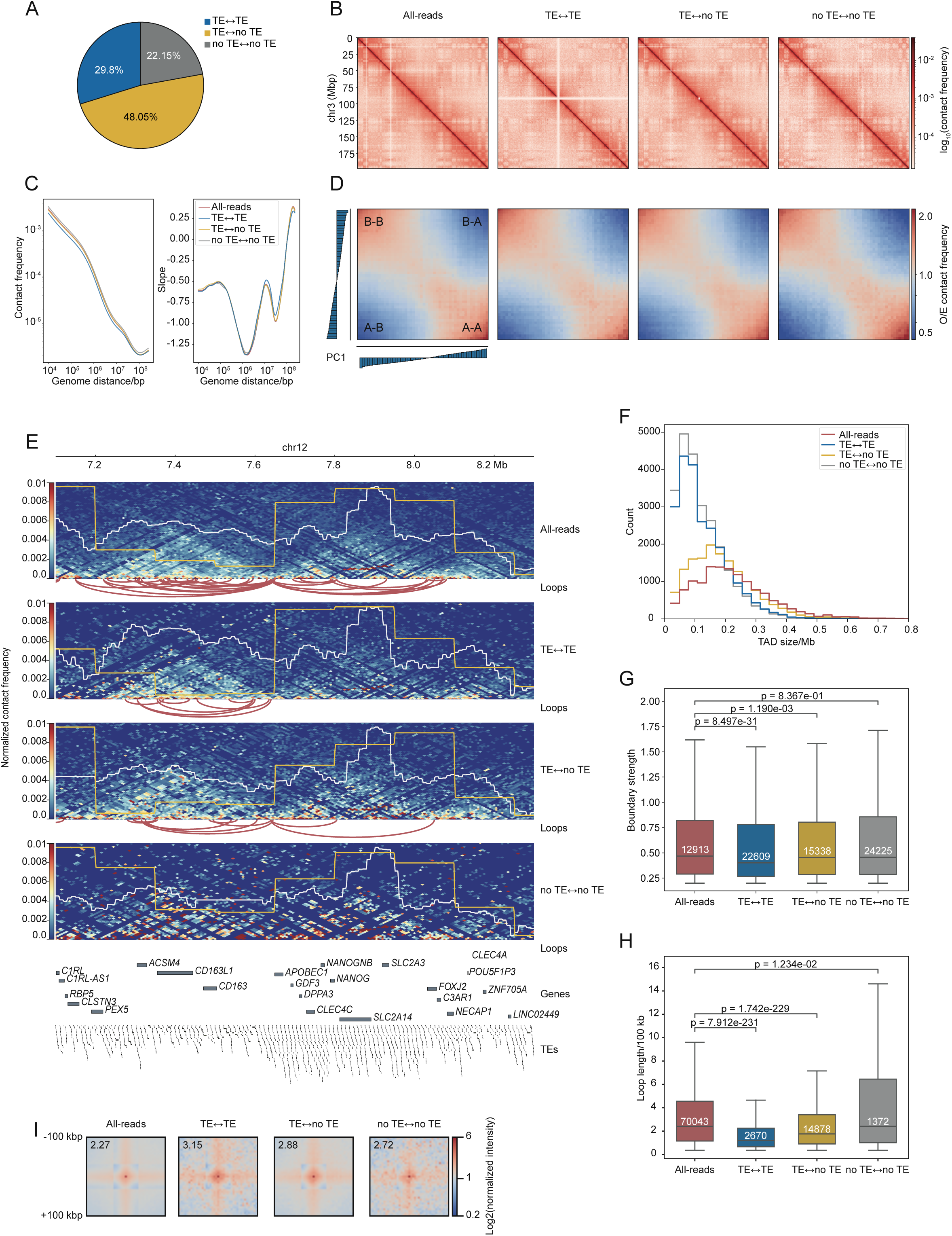
Transposable elements dominate DNA/chromatin 3D structure. (A) Pie chart showing the percentage of all reads with both ends in a TE (TE-to-TE), one end in a TE (TE-to-no TE), and with neither end in a TE (no TE-to-no TE). Hi-C data was from GSE52457 [36]. (B) Normalized contact density heatmaps for chromosome 3 using subsets of reads: (1) all-reads, (2) TE-to-TE, (3) TE-to-noTE, and (4) noTE-to-noTE matrices. (C) Plots showing contact frequency as a function of genome distance for the four matrices as in **panel B**. The left plot shows the overall contact frequency (distance in base pairs between the mapped read pairs), and the right plot shows the slopes. (D) Saddle plots showing genome-wide average O/E interaction frequency and A/B compartmentalization of the four te_hic matrices. (E) Triangle contact heatmaps at the *NANOG* locus showing the normalized contact frequency (heatmap), A/B compartments score (black line on the heatmap), insulation score (yellow line in the heatmap), and loops (red line below each heatmap) for the four matrices. The location of genes and TEs are plotted below the heatmap. (F) Histogram of TAD domain sizes in megabase pairs for the four matrices. (G) Boxplot of the boundary strength for TADs estimated for each of the four matrices. The number in white indicates the number of boundaries observed in each matrix. Significance is from a two-tailed Mann-Whitney U test. (H) Boxplots of average loop length generated from the indicated Hi-C matrices. The number of loops found in each matrix is indicated with the white numbers. Significance is from a Mann-Whitney U test. (I) Aggregate Pileup Analysis (APA) plots centered on the loop coordinate in the Hi-C matrices. The flanking 500 kbp is shown. The value in the top left corner is the score of the central pixel.

We next asked how many 3D structure elements (compartments, TADs, boundaries, loops, and contacts) could be recovered using matrices generated with or without TE reads. Comparison of the overall contact matrices at the chromosome levels revealed a similar pattern for all four matrices (**Figure 2B**). Namely, contact frequency as a function of chromosomal distance was similar for all matrices, albeit the level was reduced in the TE matrices versus the all-reads matrix (**Figure 2C, left plot**). However, the slope of all four matrices was nearly identical, indicating that the overall pattern of the four matrices was the same (**Figure 2C, right plot**). The assignment of A/B compartments was similar in all four matrices (**Figure 2D**). Saddle plots comparing all A/B compartment scores showed near-perfect significant correlations (R=∼0.98) for the all-reads matrix versus the TE matrices (**Figure 2D**).

This could be exemplified by the *NANOG/DPPA3* locus, where all four matrices were similar, except the no-TE-containing matrices were sparser (**Figure 2E**). TADs could be inferred from all four matrices, albeit some TADs were estimated to be smaller when calculated from the TE-to-TE or no-TE-to-no-TE matrices, suggesting that TADs identified using TE sequences were fragmented (**Figure 2F**). Surprisingly, boundary strength scores were significantly weaker in the TE matrices compared to the all-reads matrix, but not in the no-TE-to-no-TE matrix (**Figure 2G**). This suggests that TEs may contribute less to boundary formation. Next, we called chromatin loops. Of the total of 70,043 detected loops only 1,372 could be detected in the TE-to-TE matrix and a combined 17,548 loops in a TE-matrix (**Figure 2H**). This suggests that sub-setting the matrices by TE impairs the ability to detect loops. Curiously, the lowest number of loops was detected in the no-TE-to-no-TE matrix (1,372), although these loops were significantly longer, whilst loops detected in TE-containing matrices were significantly shorter (**Figure 2H**). Aggregate pileup analysis (APA) plots centered on the loop points in each matrix showed the highest strength in the TE-to-TE loops among all four types (**Figure 2I**). This APA suggests that TEs are a contributor to short and mid-range loop and chromatin contact formation.

### Transcription factors drive 3D genome contacts

We next sought to explore mechanisms regulating 3D structure at TEs. TFs bind to TEs [29, 37, 38], and can both make [9, 39, 40] and break [20] chromatin contacts, hence it seems likely that TFs bind to TEs to mediate 3D structure. To explore this idea, we took advantage of the large amount of TF-DNA ChIP-seq data available in mouse and human PSCs along with corresponding high-depth Hi-C data [36, 41, 42]. We collected 171 TF binding profiles (mainly from ChIP-seq data) for human and 120 datasets for mouse (**Supplementary Table 1**). Most of this data was taken from the Cistrome database [42], supplemented with the reanalysis of selected samples (Supplementary Table 1).

We devised a metric to assess if a TF was positively or negatively correlated with chromatin contacts. We did this by generating a matrix of all possible pair-wise interactions between any two loci containing the binding sites of the same TF (i.e. homotypic TF-TF contacts). We then measured the contact strength (normalized Hi-C reads) between each pair of TFs and compared these scores to random backgrounds generated from shuffled genome regions (using *bedtools shuf*) to generate a Z-score. The Z-score represents TF-TF pairs where positive scores indicate an association with increased chromatin contacts, and negative scores with decreased chromatin contacts (**Figure 3A, B, and Supplementary Table 2**). To validate this approach, we first confirmed that the Z-score did not correlate with potential confounding metrics, such as the number of TF-bound genome loci. This was the case for human (Spearman R=0.2) (**Figure 3A**) although there was a weak correlation in the mouse ChIP-seq data (**Supplementary Figure S1A**). Additionally, there was no strong correlation between the Z-score and the percentage of peaks co-bound with CTCF (**Figure 3A and Supplementary Figure S1A**), indicating that this model was not capturing a simple relationship with CTCF. We then divided the TFs into contact-formers (associated with increased chromatin contacts), and contact-breakers (associated with decreased chromatin contacts) defined using threshold Z-scores of +0.6 and −0.7 based upon the first and fourth quartiles of the Z-scores (**Figure 3A**). We termed the remaining TFs as ‘neutral’, with respect to chromatin structure. Encouragingly this approach correctly identified known contact-formers with positive Z-scores such as CTCF, RAD21, and SMC3 [39, 43, 44]. Known contact-breakers are not well described, and we did not identify the known contact-breaker ADNP (Z-score=0.06) [20], indicating there is information our model is not capturing. Overall, the model presented here suggests TFs can be divided into factors that are associated with increased chromatin contacts and those that are associated with decreased contacts.

**Figure 3.**
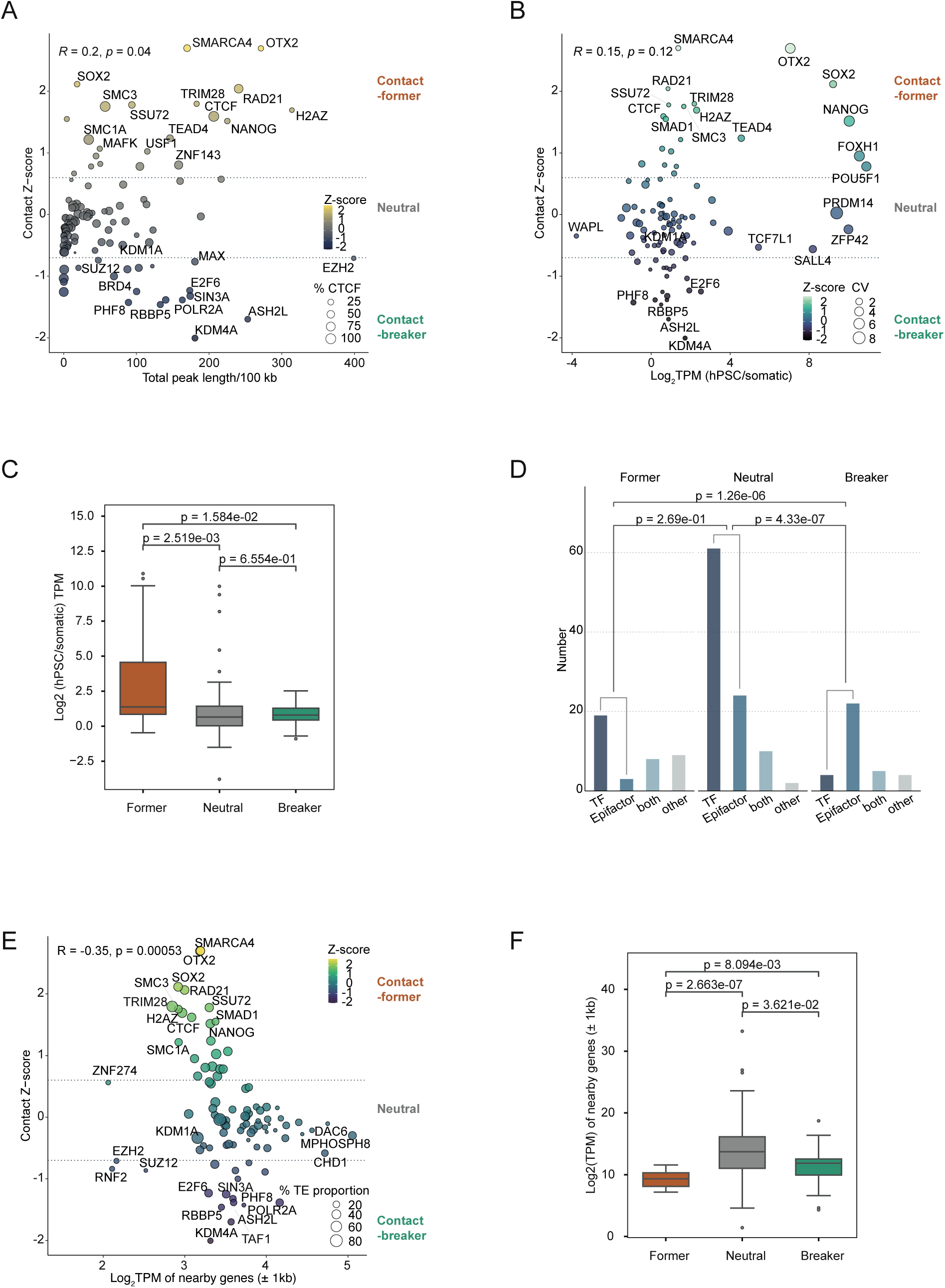
Transcription factors drive 3D genome contacts. (A) Scatter plot of contact Z-score versus normalized peak size (in kilobase pairs) for hPSCs. Only homotypic TF-TF contacts are considered. The size of the dots indicates the percentage overlap with CTCF binding sites. The colors of the dots indicate the Z-score. (B) Scatter plot of Z-score versus the fold-change expression of the indicated factors in hPSCs versus the other somatic cell types (See also **Supplementary Figure 1B**). The dot size indicates the coefficient of variance (CV) for the y-axis Log2(hPSC/somatic) expression ratio. The colors of the dots indicate the Z-score. Significance is calculated from Spearman’s R. (C) Boxplot of the TPM expression ratios for all contact-formers, breakers, and neutral factors as defined in **panel A**. Significance was from a two-sided Mann-Whitney U test. (D) Bar chart summarizing the number of TFs and epigenetic factors in the contact-former or contact-breaker categories, as defined in **panel A**. A protein was defined as a TF if it appeared in the AnimalTFDB [73], and an epigenetic factor if it is present in the Epifactor database [74]. Note that some proteins are present in both or neither database. p-values are from a Fisher exact test of TF/epigenetic factor pairs between the three contact categories, former, neutral, and breaker. (E) Scatter plot of contact Z-score versus the log2(TPM) of nearby genes with a TSS within 1 kbp 5’ or 3’ of a TF binding site. The size of the dots indicates the percentage of TF binding sites that are within a TE. The color of the dots indicates the contact Z-score. (F) Boxplot showing the expression of all genes with a TSS within 1 kbp 5’ or 3’ of a TF-bound locus, divided into formers, breakers, and neutral TFs. Significance was from a two-sided Mann-Whitney U test.

Interestingly, we noticed patterns in the predicted contact-formers and breakers. Contact-formers tended to have higher ratios of cell type-specific expression, whilst contact-breakers were expressed constitutively the same as neutral factors (**Figure 3B, C and Supplementary Figure 1B**). This is exemplified by the predicted contact-formers NANOG, SOX2, and OTX2, which are specifically expressed in hPSCs and are important for pluripotency maintenance [45]. Conversely, epigenetic factors SMARCA4, TRIM28, and H2AZ are relatively constitutively expressed (**Figure 3B and Supplementary Figure 1B**). There was also another pattern: contact-formers were significantly more likely to be TFs, while contact-breakers were more likely to be epigenetic factors (**Figure 3D**). For example, the epigenetic repressors SUZ12, EZH2, RNF2, and SIN3A, are all predicted contact-breakers. An interesting exception for epigenetic factors was POLR2A, the largest subunit protein of RNA-polymerase II, as well as BRD4, both of which are associated with actively transcribed regions and transcription start sites [46, 47], and both were predicted to be contact-breakers. Potentially POLR2A and BRD4 appear as contact-breakers as they are associated with actively transcribed regions that tend to have a looser 3D structure, and which agrees with reports that transcription disrupts local chromatin contacts at promoters [48]. Indeed, there was a weak, but significant, negative correlation (R=-0.35; p=5.3e-4) between the contact Z-score and the expression of nearby genes (**Figure 3E**). Additionally, the expression of genes with a TSS within 1 kbp of a TF-bound locus were lower for formers, compared to neutral factors (**Figure 3F**). As contact-formers are more likely to be cell type-specific factors, and cell type-specific genes also tend to have lower expression [49], this suggests they are regulating cell type-specific genes (**Figure 3C**).

### TF-TE contacts organize 3D genome structure

Next, we sought to integrate TFs, TEs, and 3D genome structure to understand if the predicted contact-formers and breakers are associated with Tes, and if their forming and breaking capability is a TE-independent function. We focused on hPSCs for this analysis. First, we calculated the proportion of TE sequence and the proportion of TE contacts at one or other end of the paired reads (**Figure 4A and Supplementary Figure 2A**) in the TF-bound Hi-C read pairs versus the contact Z-score (**Figure 4A**). Interestingly, predicted contact-former TFs tended to be bound to TEs (**Figure 3E, 4A**), and have a higher frequency of contacts involving TEs, compared to the neutral or contact-breaker TFs (**Figure 4A, Supplementary Figure 2A**).

**Figure 4.**
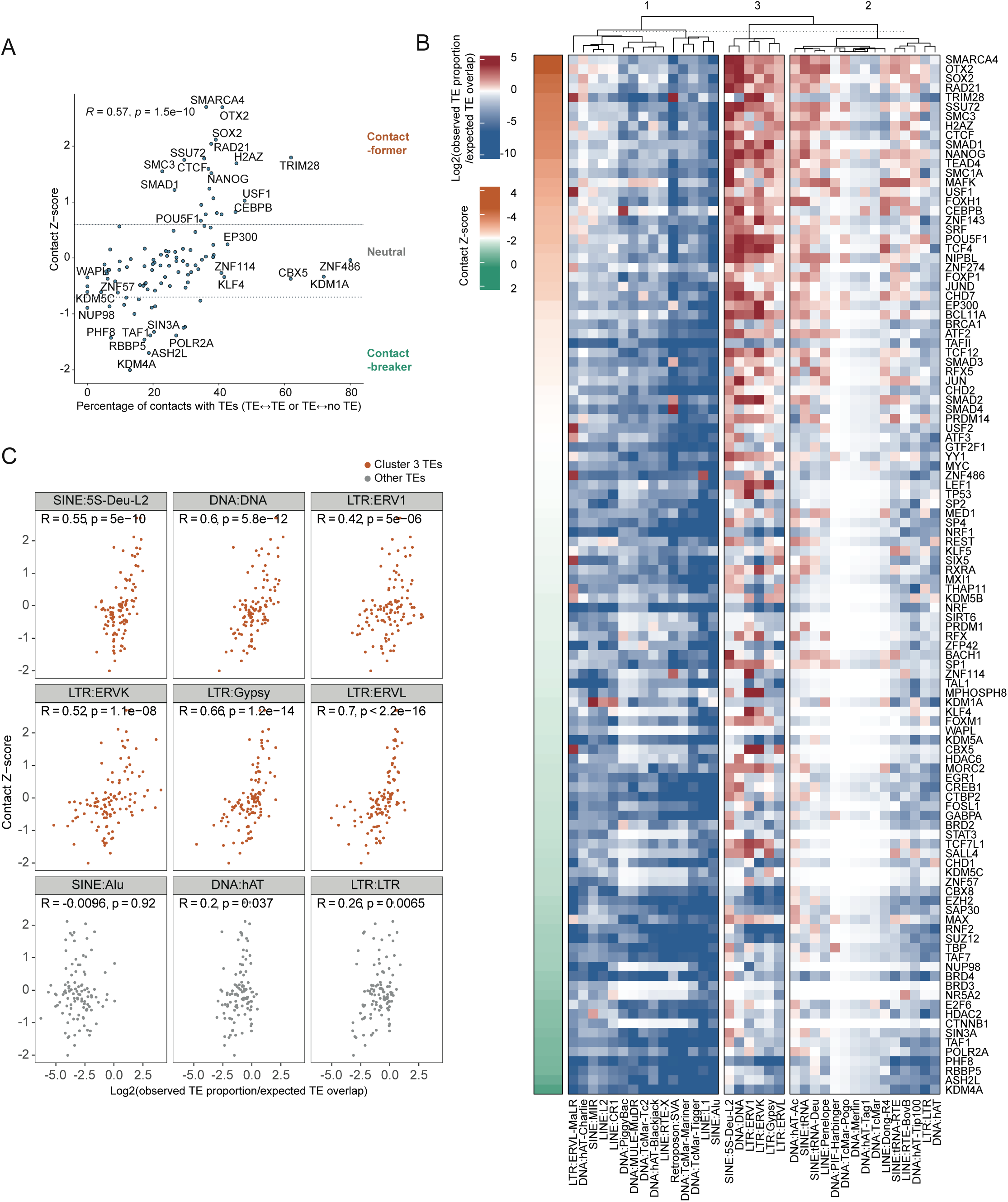
TF-TE contacts drive 3D genome structure. (A) Scatter plot showing the correlation between the contact Z-score versus the TE proportion, defined as the percentage of HiC read pairs with one or more reads inside a TE sequence, and one or more read within 100 bp of the indicated TF binding site (See also **Supplementary Figure 2A**). (B) Heatmap of TE enrichment TF binding loci. TE proportions were calculated for each TE and TF, and the TE proportions were further normalized to the TE proportion expected by chance alone. The binding sites of the TFs were compared to an expected (randomized shuffled) background of all TEs. The resulting heatmap was clustered by *k*-mean clustering (*k* = 3). The TFs are ordered by their average contact Z-score from high to low versus the log2 fold-change of observed TE proportion/expected TE overlap. (C) Scatter plots of selected TE types showing the contact Z-score versus the log2 fold-change of observed TE proportion against expected TE overlap, which represents the enrichment of TE sequences for each TF (as in **panel B**). Each dot is the binding data for a transcription/epigenetic factor from **panel B**. The top two rows are from Cluster 3 in **panel B**, and the bottom row is a selection of TEs from Cluster 1 and Cluster 2. The Pearson R correlation is shown, along with the p-value.

To understand the TE types that the TFs are binding to, we correlated the Z-score against TE proportion in TF binding sites (**Figure 4A and Supplementary Figure 2A**). As TEs occupy a substantial fraction of the genome, this proportion was normalized by comparing the observed TE percentage versus a (shuffled) expected background of TFs and measuring the observed/expected TE overlap by chance alone (**Figure 4B**). Interestingly, the heatmap suggests that TEs involved in contact-forming include LTRs (ERV1, ERVL, and ERVK), DNA, and SINEs:5S-families of TEs (**Figure 4B, group 3**). Unexpectedly, TE overlaps were depleted at contact breakers (**Figure 4A, B**). This suggests that contact-formers are mainly binding to TE sequences to form chromatin-contacts, however, our predicted contact-breakers appear to avoid TEs. Subplots of the correlations of individual TFs versus TE enrichment showed clear correlations between Z-score and enrichment of TE-mediated contacts, supporting the idea that contact-formers (high-Z-scores) were more likely to bind TEs whilst contact-breakers were less likely to bind TEs (**Figure 4C**). This effect was specific for cluster 3 TF/TEs, as clusters 1 and cluster 2 had poor or no significant correlation between the contact Z-score and TE enrichment, for example, SINE:Alu (**Figure 4C**). Overall, our computations can divide TFs into predicted contact-formers and -breakers of 3D genome structure, and contact-formers are associated with TE-driven chromatin contacts that link specific TE types.

### Reduced expression of predicted contact-formers *SMARCA4* and *MAFK* disrupts 3D chromatin structure at TEs

Our approach predicts potential contact-formers and contact-breakers among the TFs at TE-bound loci, so we set out to experimentally validate candidates for those roles. We selected two predicted contact-formers at TEs: the epigenetic factor SMARCA4 (BRG1) and the TF MAFK. SMARCA4 is a member of the BAF complex involved in chromatin remodeling [50], which has important roles in epigenetic control by modulating nucleosome positioning. MAFK is a BZIP-family transcriptional repressor [51]. Neither of these two factors has been reported to have a role in regulating pluripotency in hPSCs.

Knockdown (KD) of both *SMARCA4* and *MAFK* in hPSCs achieved between 60-90% KD (**Supplementary Figure 3A**). Bulk RNA-seq of the KDs suggested only small changes in overall genes expression (**Supplementary Figure 3B**), however, the *SMARCA4* KD caused a notable change (**Supplementary Figure 3C**), with 273 and 408 significantly up- and down-regulated genes (**Supplementary Figure 3D, E**). We performed Hi-C to explore the changes in the 3D genome structure and applied our te_hic pipeline. Quality control metrics were similar for the two KDs (**Supplementary Figure 4A-D**), and, gross Hi-C matrices were unchanged, A/B compartment changes were relatively minor, and TADs were unaffected (**Supplementary Figure 4E-H**). However, the differential contact frequency as a function of distance revealed that the knockdown of *SMARCA4* caused a large drop in contact frequency at short ranges, (**Figure 5A-C**). Chromatin loop number and loop lengths were decreased in the KD samples (**Supplementary Figure 4I**). Pileups of sh*Luc* loops from the four te_hic matrices against the four matrices of both control and KDs showed a decrease of the central pixels in TE loops of KDs, indicating that KD of SMARCA4 and MAFK disrupted chromatin loops, and this occurred at TE-anchored loops (**Figure 5D**).

**Figure 5.**
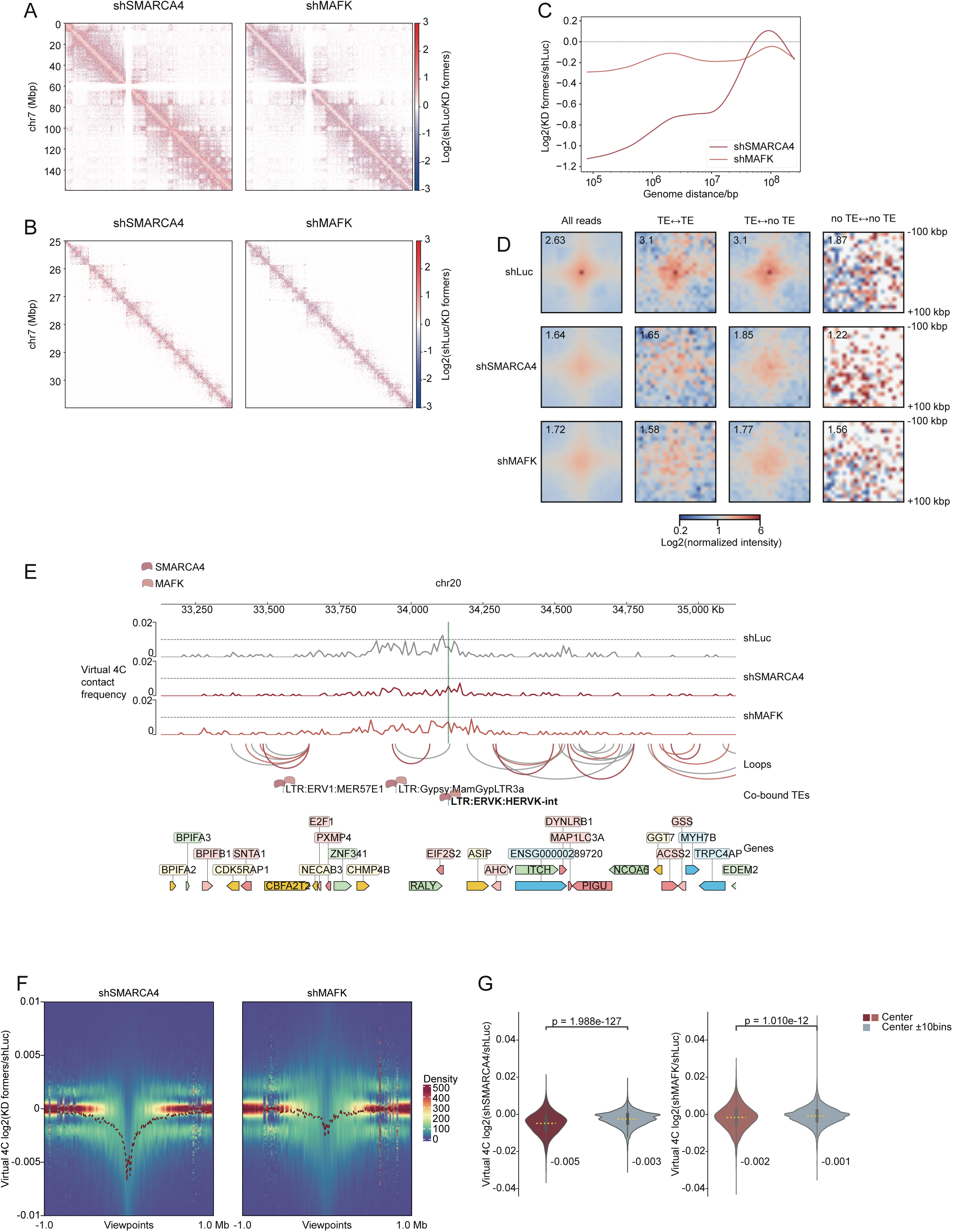
Reduced expression of contact formers leads to disrupted chromatin interactions at TEs. (A) Differential contact heatmaps of sh*Luc* transfected hPSCs versus hPSCs transfected with shRNAs targeting the predicted contact-formers *SMARCA4* and *MAFK*. The heatmaps show all of chromosome 7 at 150 kb resolution. (B) Differential heatmaps of a zoomed-in region (chr7:25000000-31000000) at 10 kb resolution. (C) Line plot showing the differential contact frequency as a function of distance relative to the sh*Luc* data. (D) Pileups of loops called from the four te_hic matrices of shLuc relative to the four matrices of sh*Luc*, sh*SMARCA4*, and sh*MAFK* KD Hi-C data. (E) Virtual 4C view with a cluster 3 TE (LTR:ERVK:LTR5_Hs) as the viewpoint showing the same region of the genome (chromosome 4) for sh*Luc*, sh*SMARCA4*, and sh*MAFK* Hi-C data. SMARCA4 and MAFK are bound at the center point. (F) Density heatmap of log2 comparison between knockdowns and the control of virtual 4C values with cluster 3 TEs bound by SMARCA4 or MAFK as viewpoint centers and the flanking ± 1 Mbp (left: sh*SMARCA4*, right: sh*MAFK*). (G) Violin plots of log2 comparison between knockdowns and the control of virtual 4C values with cluster 3 TEs bound by SMARCA4 and MAFK as viewpoints in the center and the flanking ± 100 kbp (left: sh*SMARCA4*, right: sh*MAFK*). Significance was from a two-sided Mann-Whitney U test.

In addition, we assessed the effect of the KDs on loci rich in TE sequences. We generated virtual 4C plots from the Hi-C data using CoolBox [52]. In virtual 4C a specific genomic region serves as a central viewpoint, and interactions are measured relative to the viewpoint. Virtual 4C using cluster 3 TEs bound by SMARCA4 or MAFK as the viewpoint center showed a decrease in contact frequency for both the *SMARCA4* and *MAFK* KDs, compared to the sh*Luc* control (**Figure 5E, F and Supplementary Figure 5A-C**). Aggregate virtual 4C plots centered on cluster 3 TEs illustrate this decline for both *MAFK* and *SMARCA4* KDs (**Figure 5F**), with the largest decline seen in the *SMARCA4* KD. Quantitation of these effects showed a significant decrease in chromatin contacts originating at cluster 3 TEs for both SMARCA4 and MAFK KDs (**Figure 5G**). Based on these results, we conclude that SMARCA4 and, to a lesser extent, MAFK, build chromatin contacts at TEs, as disrupting them reduced chromatin contacts originating at TEs and disrupted TE-mediated chromatin loops.

## Discussion

The interphase nucleus is highly structured at multiple scales, from megabase domains down to individual nucleosome structures. However, the determining features of this 3D structure remain unclear. Here, we show that TEs make a major contribution to 3D genome structure, mediated by TE-bound factors that build chromatin contacts mainly at the scale of chromatin loops and short-range chromatin contacts. TE sequences are permanently integrated into the genome and are relatively invariant as most TEs in human and mouse are no longer capable of transposition. This would help explain why Hi-C data is different between species, but similar between cell types of the same species [53, 54]. The distribution and constituent TEs are divergent between species but are essentially identical between cell types. Interestingly, Hi-C experimental data and chromatin contact perturbations can be computationally predicted using machine-learning approaches from DNA sequence alone [55, 56], suggesting that 3D structure is indeed encoded in the linear sequence. As TE sequences comprise nearly half of the genome, potentially Hi-C data could be predicted from TE sequences alone.

Experimental exploration of the roles of TEs in 3D structure has been limited to date, as the massive amount of TE sequences in the genome makes designing experiments challenging. A novel experimental technique based on Hi-C (4Ttran-seq), that specifically targets TE-to-TE 3D genome interactions, identified long terminal repeats (LTRs) bound by TFs as mediators of chromatin contacts in A/B compartments and TADs [30]. The knockout of specific HERVHs on chromosomes 4 and 13 disrupted TAD domain boundaries [22]. Conversely, knockouts of an L1M3f (chromosome 10) or LTR41 (chromosome 8) caused the fusion of TADs [54], albeit both TEs contain CTCF binding sites which might explain the TAD fusion. In our experiments knocking down *SMARCA4* and *MAFK,* we did not observe TAD disruptions, instead we only saw changes in TE-mediated chromatin contacts, suggesting multiple mechanisms to regulate 3D structure.

Phase separation may also contribute to TE drive 3D conformational changes. LTR sequences, when transcribed, can influence local transcriptional condensates [57], and disruption of phase separation using small molecules can radically alter the 3D genome structure [58]. Potentially, there are deeper links, considering many TEs are often expressed as parts of long non-coding RNAs [9], that localize on chromatin [59], to drive phase separation and affect 3D genome structure [57, 60]. Potentially, in addition to TFs, TE-containing non-coding RNAs interact directly with their DNA sequences to modulate chromatin interactions [61, 62]. TE sequences are often in heterochromatic regions, which are found at the nuclear periphery. This is especially the case for LINE L1 elements, which are associated with laminin-associated domains [63]. There is a link between TEs and the formation of chromatin condensates along with euchromatin and heterochromatin formation and modulation.

### Conclusions

While TEs have been known to modulate TAD and 3D genome formation, a comprehensive understanding of how TEs modulate 3D structure would enable the use of TEs as a tool to reengineer 3D genome arrangements [27, 64]. Our data suggests that the Hi-C signal is composed of around 48% of TE-to-TE contacts and a further 22% between TE-to-not-TE loci (**Figure 2A**). These data indicate that TE sequences constitute around ∼70% of the 3D signal detected in typical Hi-C experiments. Potentially over evolutionary time, TE insertions are responsible for modulating 3D genome structure and so generating new gene regulatory networks. We believe our tool, te_hic, could be easily expanded with the addition of new species to explore TE contributions to 3D structure during evolution, and how those structures are reshaped in different species. Overall, our data suggests that TEs are responsible for a majority of the 3D genome structure in mammalian cells. This mode is driven by TFs that are recruited to TE sequences that bring regions of DNA into proximity.

## Methods

### Analysis of Hi-C data

Reads were mapped to the human reference genome (hg38) using bowtie2 with the settings ‘--mm --very-sensitive’ [33] and processed with te_hic with default parameters and HiC-Pro pipelines [32]. The resulting matrices were normalized by iterative correction (IC) matrix balancing with cooler [65]. Different levels of the 3D structure were identified with cooltools [66]. Compartments were defined at 150 kb resolution with GC content as a phasing track, and the Eigenvector of the first principal component (PC1) was used to indicate the compartments where regions with negative values were compartment B and ones with positive values were compartment A. TAD boundaries were called at 10 kb resolution, and TAD domains were defined from the combinations of all the boundaries within the same chromosome with a maximum length of 1.5 Mb. Loops were called using Peakachu [67] with pre-trained models and a threshold of 0.8 at 5 kb resolution for Hi-C data in GSE52457 [36] and 10 kb for sh*Luc*, sh*SMARCA4* and sh*MAFK* data. Virtual 4C was performed using CoolBox [52] at the center loci of TEs spanning to a 1 Mbp window. Retrieved matrices were used to generate aggregate virtual 4C density heatmap using ComplexHeatmap R package [68]. Contact frequency as a function of genomic distance and its slope and saddle plots were performed with default parameters using cooltools. Pileup analysis was performed using loops called by Peakachu using ICE-balanced Hi-C matrices with coolpup.py [69]. Tracks were generated with pyGenomeTracks [70].

### Selection and analysis of ChIP-seq data

For human, a total of 171 hPSC ChIP-seq data sets for 108 transcription factors/other DNA-associated proteins were collected. Out of 171 samples, 106 and 63 were downloaded from the Cistrome Data Browser and the ENCODE Consortium portal, respectively. Additionally, ChIP-seq for THAP11 (accession number: GSM1505796 [71]) and WAPL (accession number: GSM2816640 [72]) were obtained from the GEO database in raw SRA format and reanalyzed. For mouse, samples with less than 5000 peaks were removed from the analysis. TFs were annotated using the AnimalTFDB [73], and epigenetic factors were defined using the Epifactors database [74].

### TF-bound Hi-C pairs and contact Z-scores

TF-specific chromatin contacts were obtained using a computational utility ‘getContacts’ from te_hic pipeline. ChIP-seq narrow peaks and Hi-C read pair BEDPE file from the all-reads matrix generated by te_hic were used as the inputs. For each TF, all possible pair-wise interactions for the same TF at different loci in Hi-C pairs were counted. The number of Hi-C reads in a 10 kb window centered on each pair of TF pair-wise binding was collected. This value as compared to an ‘expected’ random null background for each TF-specific pairs which was generated by shuffling ChIP-seq peaks by ‘bedtools shuf’ command for 10 times for each ChIP-seq data [75]. The number of observed pairs was normalized by subtracting the number of randomly expected pairs and dividing by the sum of peak length (in base pairs) to generate a Z-score for each TF which we designated as the contact Z-score. The normalized number of Hi-C pairs for each TF was defined as:

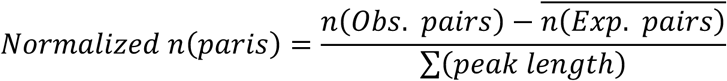

### Analysis of gene expression in hPSCs, somatic cells, and tissue

Preprocessed and quantified RNA-seq data at the transcript level of hPSCs, somatic cells, and tissue samples were obtained from [9]. TPM (transcript per million) values less than 0.01 were corrected to 0.01 to avoid producing infinity or extreme values. TPM for different transcripts of the same genes was averaged based on GENCODE v32 annotation [76]. Coefficient of variation (CV) scores per gene were calculated to estimate the cell type-specificity of the genes.

### TE enrichment in ChIP-seq peaks

The RepeatMasker track for the hg38 (GRCh38.p3) assembly from the UCSC genome browser was used for TE annotation. Repeats marked with a ‘?’ in ‘repClass’ or ‘repFamily’ columns were excluded as unclear classifications. Repeats such as low complexity, simple repeat, satellite, and RNA and TEs that mapped to mitochondrial chromosome and position-unknown contigs or patch scaffolds were also removed in this study.

The overlap between ChIP-seq peaks and TEs reported in RepeatMasker track was determined using ‘bedtools intersect’ command. To assess whether the observed inter-section between ChIP-seq peaks and TE loci in the genome was due to random chances, 1,000 shuffled sets were generated for each ChIP-seq dataset (peaks resized to 100 bp using the genomicRanges R package [77]) with ‘bedtools shuffle’ command. The shuffled data sets were then used to intersect with TEs and serve as an empirical or null background. For each TE, the expected random intersection with the shuffled data sets was determined as the mean over 1,000 permuted sets. Finally, the observed overlaps between ChIP-seq peaks and TEs was compared to the random expected overlaps between shuffled peaks and TEs. Log2 fold-change of observed TE proportion/expected TE overlap was calculated by the following equation:

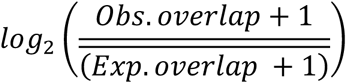

### TE proportion in TF-TF contacts

Resized ChIP-seq peaks were used to intersect the TF-bound Hi-C pairs described above to generate ‘core contacts’. A utility from te_hic pipeline, ‘assign_te.py’, was then used to assign TEs to the interacting loci of re-sized contacts. The proportion of whether containing TEs or not of all contacts for a TF was then calculated.

### Cell culture and knockdown in hPSCs

HEK293T cells were cultured in DMEM high-glucose media (Gibco, C11995500BT) with 10% (v/v) FBS (Gibco, 10091148). hPSCs were maintained on plates coated with Matrigel (BioCoat, 356231) in mTESR1 medium (Stem Cell Technologies, 85850). All cells were cultured in an incubator at 37 °C and 5% CO_2_ for the entire process. Fresh mTESR1 medium was fed daily until the cells reached a confluency of 80–90%. In general, the cells were dissociated using Accutase (Sigma Aldrich, A6964) and passaged at a 1:6 ratio in 3–4 days. For hPSCs, Y-27632 (MedChemExpress, HY-10583) was added to the medium at each passage.

shRNA sequences were synthesized and inserted into a pLKO.1 vector. shRNA sequences used: sh*MAFK:* 5’-CGATGATGAGCTGGTGTCCATGTCA-3’, sh*SMARCA4:* 5’-CCGAGGTCTGATAGTGAAGAA-3’, sh*LUC*: 5’-CGTACGCGGAATACTTCGA-3’.

Lentivirus expressing shRNAs down-regulating target genes were packaged using HEK293T in a third-generation lentivirus packaging system. The cells were co-transfected with a mixture of three vectors: psPAX, pMD2.G, and plasmid of shRNA. Lipofectamine 3000 (Invitrogen, L3000015) was used for the virus package process according to the manufacturer’s instructions. Lentivirus was infected into 200,000 hPSCs in a 10 cm dish. The medium was changed 10 hours after infection to fresh mTESR1 medium. Puromycin (1μg/mL) (Selleck, S7417) was added to select cells for a further 72 hours. RT-qPCR was performed to validate the knockdowns. qPCR primers used were: *MAFK*-F: 5’-CGACTAATCCCAAACCGAAT-3’, *MAFK*-R: 5’-ACATGGACACCAGCTCATCA-3’, *SMARCA4*-F: 5’-CAGAACGCACAGACCTTCAA-3’, *SMARCA4*-R: 5’-TCACTCTCCTCGCCTTCACT-3’, *ACTB*-F: 5’-CATGTACGTTGCTATCCAGGC-3’, *ACTB*-R: 5’-CTCCTTAATGTCACGCACGAT-3’.

### RNA-seq and data analysis

RNA was extracted by using RNAzol® RT (MRC, RN190-200) following the manufacturer’s instructions. Samples were prepared for sequencing with the RNA-seq NEB Next Ultra RNA Library Prep Kit (7530, NEB). Samples were sequenced on an Illumina Novaseq 6000 platform.

RNA-seq data was analyzed as previously described [78], except GENCODE v42 was used. Differential expression was determined using DESeq2, and a gene was considered differentially expressed if it had a fold-change of at least 1.5-fold and a Bonferroni-Hochberg corrected p-value less than 0.01. All other analysis was performed using glbase3 [79]. All RNA-seq experiments were performed in biological duplicates.

### Hi-C in hPSCs

Cells were transfected in two 10 cm dishes for each target gene. Cells were selected with puromycin and harvested at around 80% confluency after infection. Each sample had at least 8×10^6^ cells, at which point Hi-C sample processing was performed. All Hi-C experiments were performed in biological duplicate. The cells were cross-linked with formaldehyde. The culture medium was discarded and 25 ml of serum-free DMEM medium containing 2% formaldehyde was added to each dish. Cells were cross-linked at room temperature for 10 mins. At 2.5 min, 5 min, and 7.5 min the dish was slightly shaken to ensure adequate cross-linking. After 10 min of cross-linking, 2.5 ml of 2.5 M glycine solution was added, and the dish was gently mixed to ensure that formaldehyde and glycine were fully mixed. Cells were incubated at room temperature for 5 minutes. After neutralizing the dish was transferred to ice. Using a disposable cell scraper, the adherent cells were scraped off and transferred into a 50 ml centrifuge tube. Cells were centrifuged at 800g at 4℃ for 10 mins. After centrifugation, the supernatant was discarded. 1 ml of ice-cold PBS was added to fully suspend the cells and they were transferred into 1.5 ml centrifuge tubes. 10 ul of the cell suspensions was used to count the number of cells, and ensure that each sample contained at least 5×10^6^ cells. The remaining cell suspension was centrifuged at 800g at 4℃ for 10 mins. After centrifugation, a pipette was used to remove the supernatant and remove all the liquid as far as possible without disturbing the cell pellet. Processed samples were frozen in liquid nitrogen and then stored at −80℃. After formaldehyde cross-linking fixation, the proteins involved in chromatin interactions in the genome were frozen and then subjected to Hi-C as described in [80].

## Declarations

### Ethics approval and consent to participate

Not applicable

### Consent for publication

Not applicable

### Raw data accessions

The sequencing data that contributes to this study was deposited in GEO with the accession number GSE248863. The te_hic software is available online at: https://github.com/oaxiom/te_hic. Other materials are available on request to the corresponding author.

### Authors’ contributions

LY. S. performed the majority of the bioinformatic analysis, with assistance from I.A.B., XM.Z., ZQ.H. and A.P.H. Z.X. performed the experiments, with the assistance of G.M., XL. F. and S.L. A.P.H., LY.S. and Z.X prepared the manuscript. JP.H., A.S. and A.P.H. revised and edited the manuscript with the assistance of all authors. XM.Z acquired funding. A.P.H designed the study, acquired funding and supervised the project.

## Acknowledgments

We thank Dr. Jacek Majewski of McGill University for discussions and support.

