## Supplementary Figures for "Dissecting the Contribution of Transposable Elements to Interphase Chromosome Structure"

Supplementary Material

**Supplementary Table 1 – Summary of the ChIP-seq data of human and mouse PSCs used in this study.**

**Supplementary Table 2 – Predicted contact forming/breaking potential of TFs in hPSCs and mPSCs.**

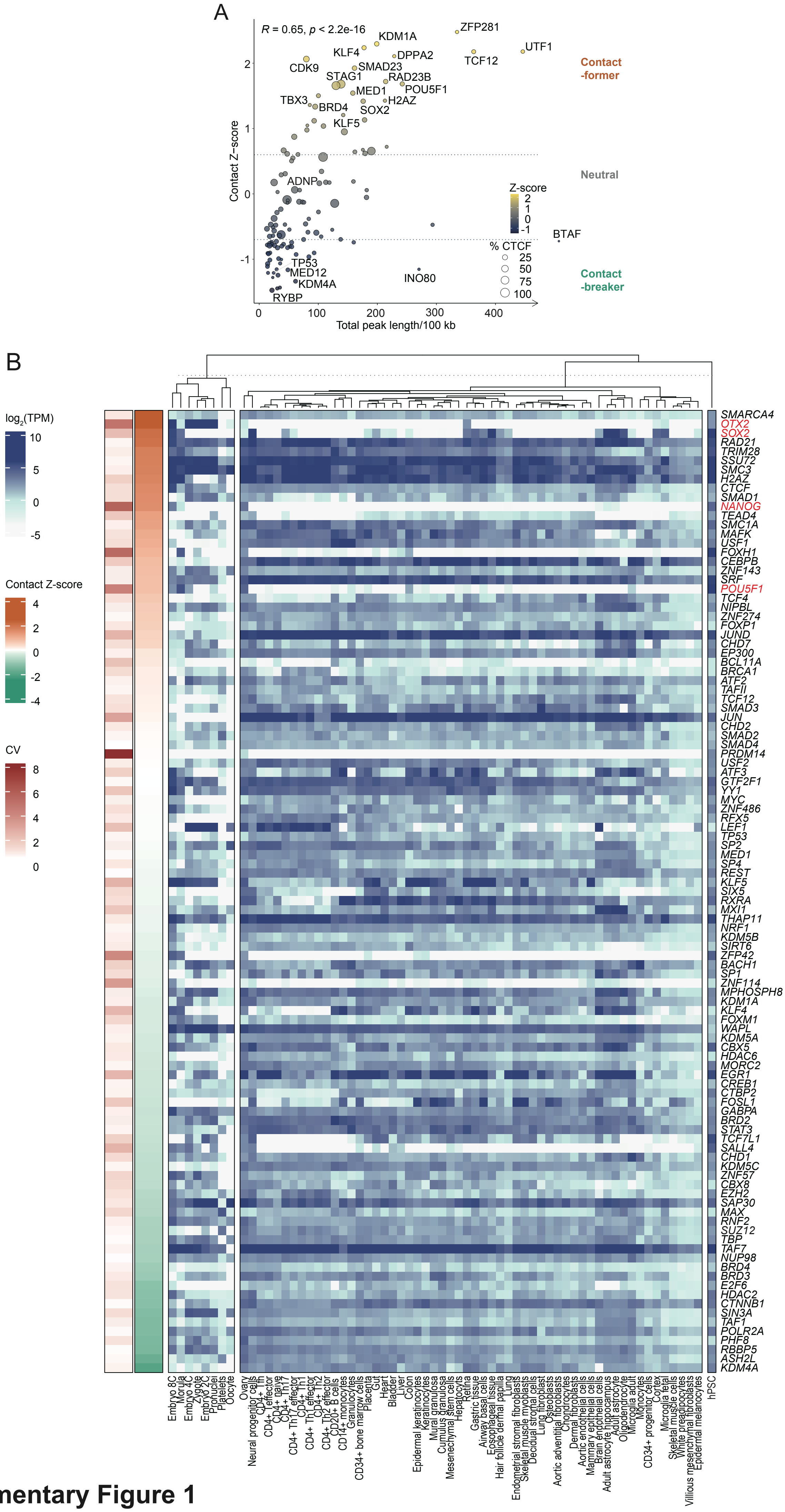


**Supplementary Figure 1**

**Supplementary Figure 1. Contact Z-score in mouse pluripotent stem cells and expression specificity of contact-formers**

(A) Scatter plot of contact Z-score versus normalized peak size (in base pairs) for mouse PSCs. Only homotypic TF-TF contacts are considered. The size of the dots indicates the percentage overlap with CTCF binding sites. Hi-C data was from GSM4074304 (Justice et al, 2020) and GSM4647542 (Carico et al, 2021).

(B) Heatmap of expression for the genes for the TFs and epigenetic factors explored in this study. Expression was measured in hPSCs versus a panel of somatic cell types. Rows of the heatmap is based on their contact Z-score from **Figure 3A**. RNA-seq data is described in (Babarinde et al, 2021).

A

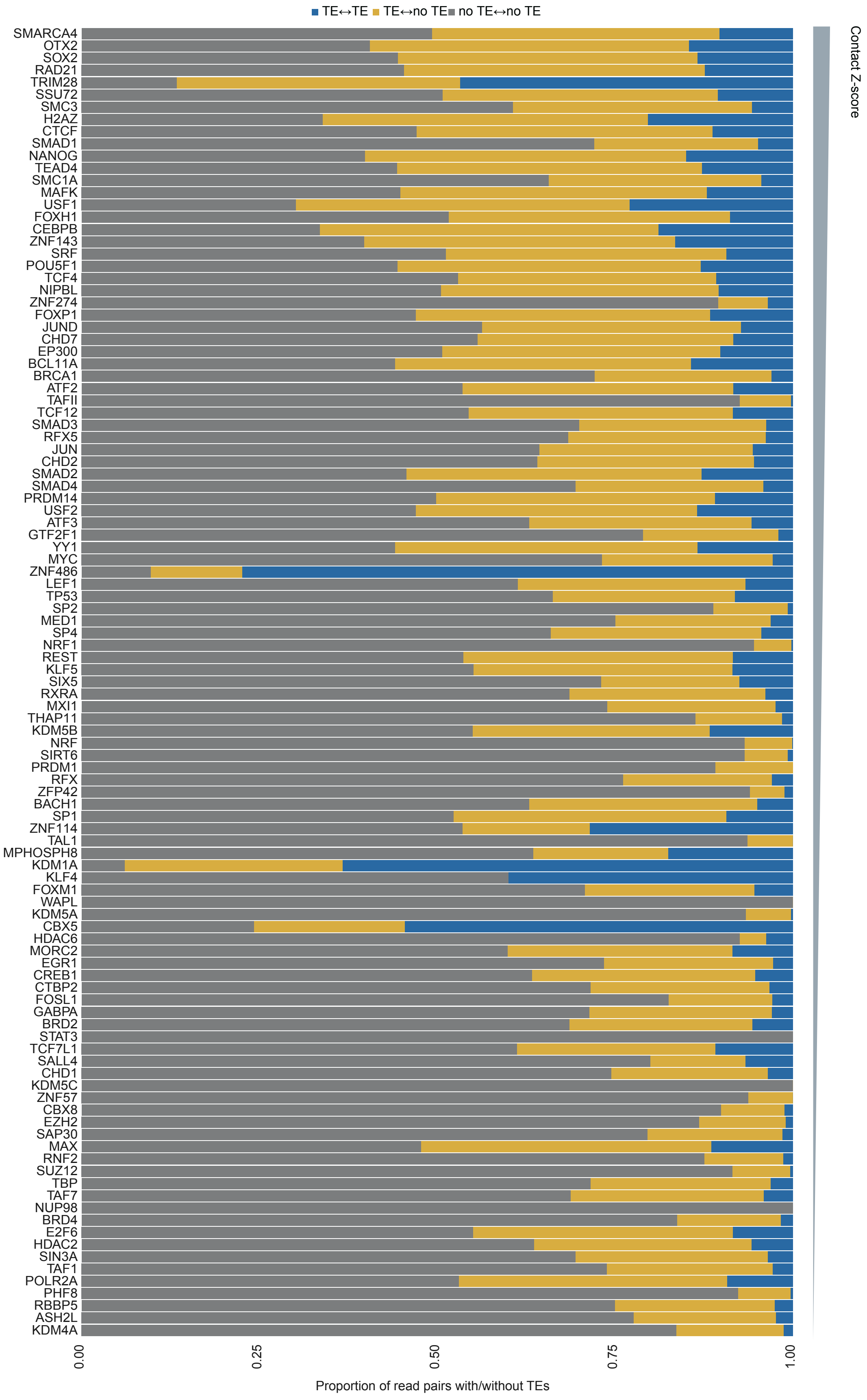


### Supplementary Figure 2

**Supplementary Figure 2. Proportion of TF-bound Hi-C read pairs overlap with TEs**

(A) Cumulative bar plot of the proportion of TEs at one or both ends of each TF-specific contact, bars are ranked by the contact Z-score from **Figure 3A**.

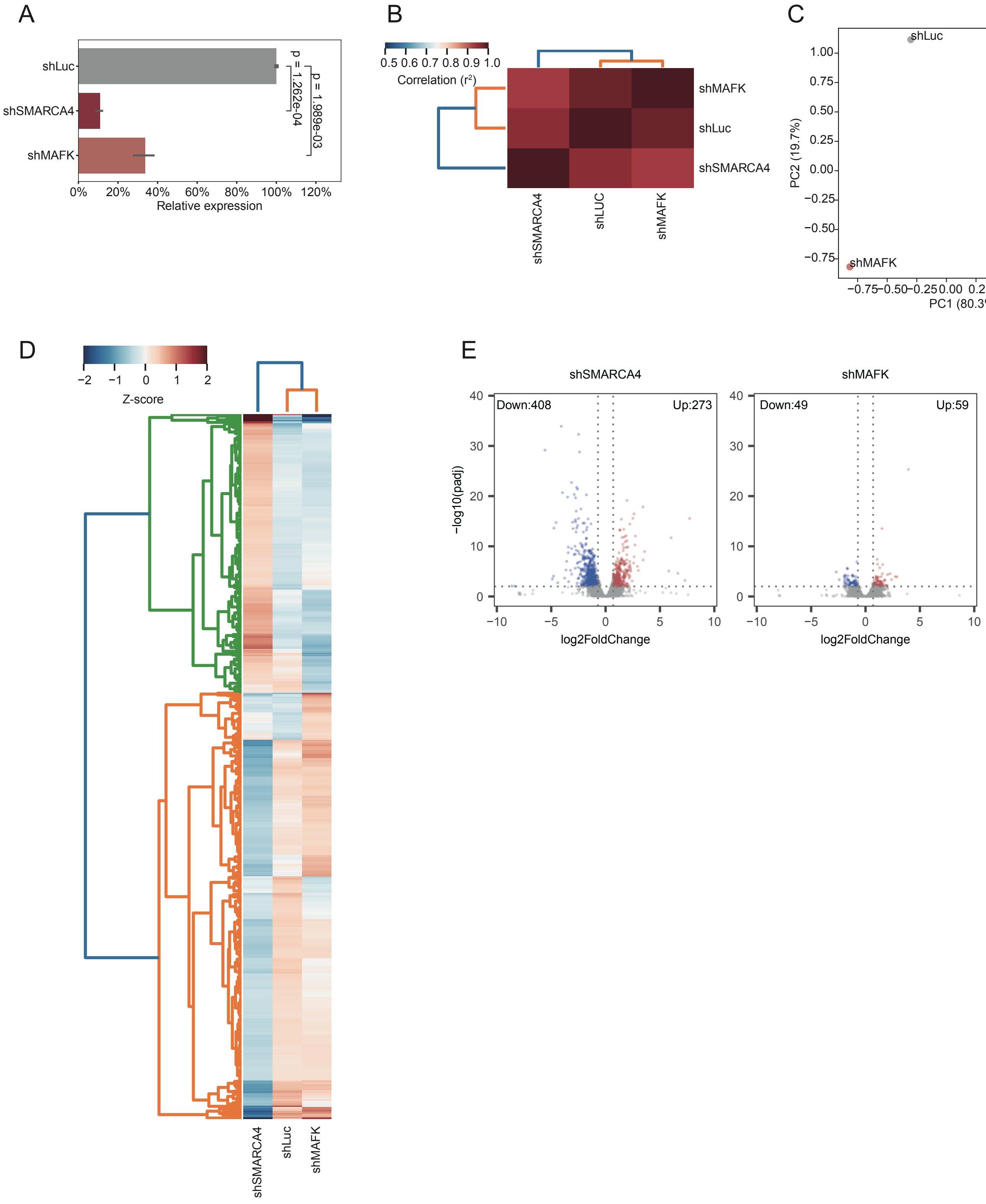


**Supplementary Figure 3**

**Supplementary Figure 3. Knockdown of candidate contact-formers *SMARCA4* and *MAFK*.**

(A) RT-qPCR for the indicated genes in the indicated knockdowns. Statistical significance is from a two-tailed unpaired Student's t-test.

(B) Coefficient of determination ($R^2$) of merged RNA-seq samples.

(C) Principal component analysis (PCA) of merged RNA-seq samples.

(D) Heatmap of all differentially expressed genes in the indicated knockdown.

(E) Volcano plots for all differentially expressed genes in shS*MARCA4* and sh*MAFK* versus sh*Luc* control. A gene was considered differentially expressed if it had a fold-change of at least 2-fold and a Bonferroni-Hochberg corrected p-value (padj) less than 0.01.

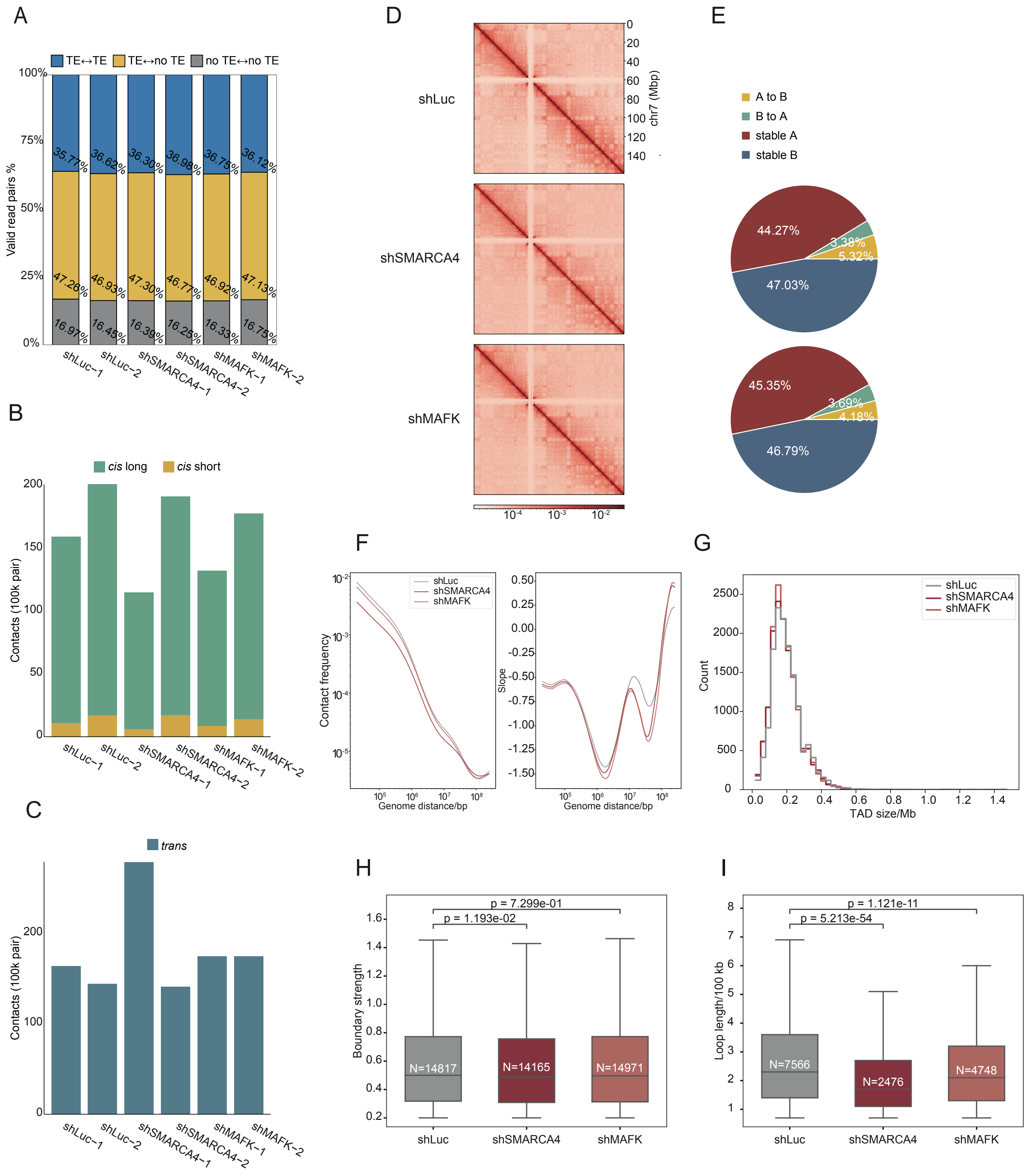


### Supplementary Figure 4

**Supplementary Figure 4. Hi-C metrics for the knockdown of the predicted contact-formers *SMARCA4* and *MAFK***

(A) Cumulative bar chart showing the proportion of reads assigned to each matrix by te_hic.

(B) Number of detected chromatin contacts in the indicated Hi-C knockdown data. *cis*-short was defined as within 20 kbp, and *cis*-long was defined as over 20 kbp.

(C) Bar chart showing the number of *trans* contacts in the indicated knockdown.

(D) Heatmaps of the normalized contact intensity of chromosome 7 for the indicated knockdowns using the all-read matrices generated by te_hic.

(E) Pie charts showing the proportion of compartment switches in the indicated knockdowns, relative to the sh*Luc* control knockdown. Compartments were defined at 150 kb resolution. Only bins with same sign in PC1 values in two replicates were kept.

(F) Contact frequency against genomic distance. Right panel: slopes of the curves.

(G) Histogram of the TAD domain size distribution in the indicated knockdowns.

(H) Boxplot showing the TAD boundary strength, and the number of boundaries detected (white). Statistical significance is from a two-tailed Mann-Whitney U test.

(I) Boxplots of loop length of the indicated knockdowns. Number of loops are indicated (white). Statistical significance is from a Mann-Whitney U test.

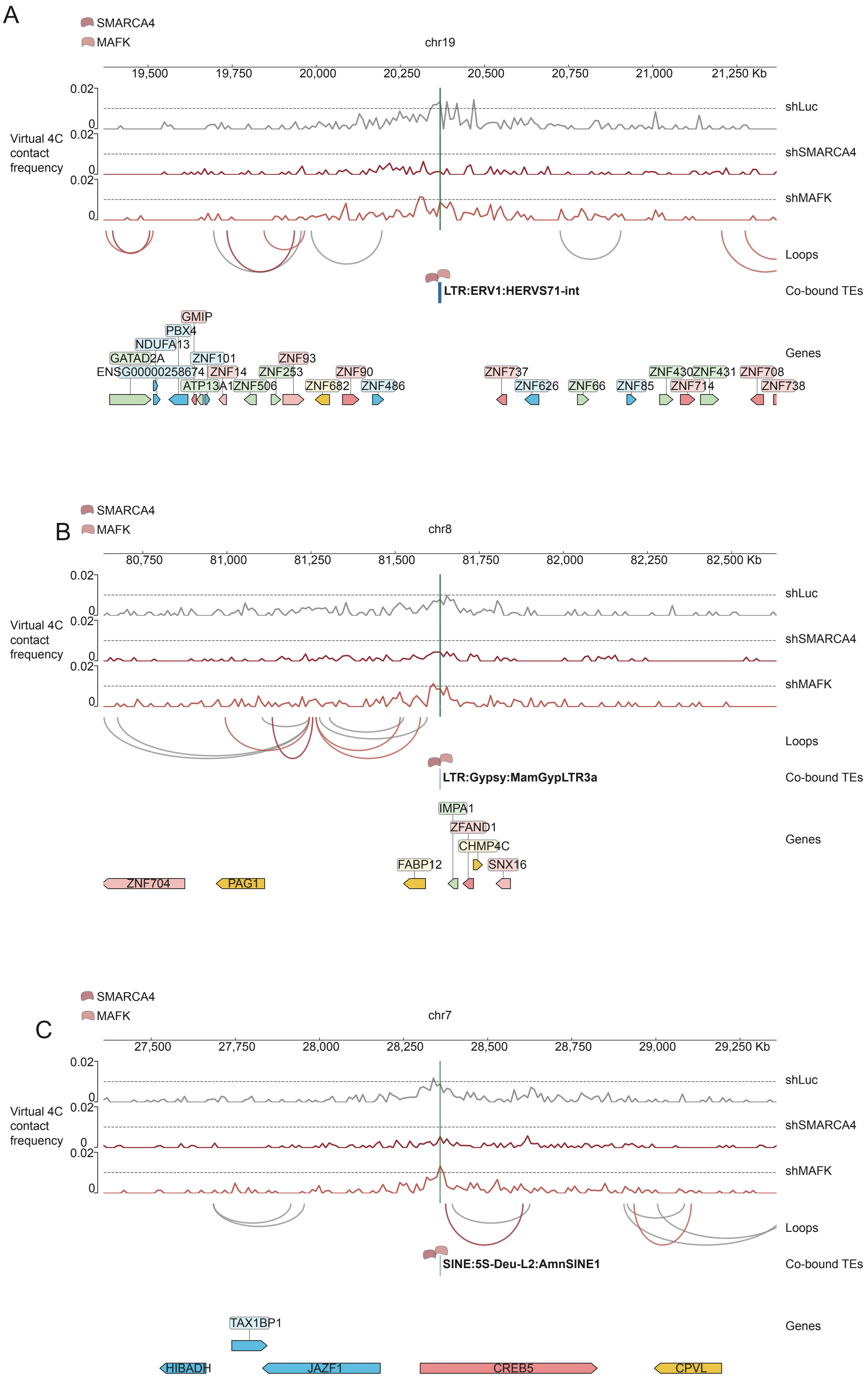


### Supplementary Figure 5

**Supplementary Figure 5. Virtual 4C views at selected loci for the contact-formers.**

(A) Virtual 4C views of sh*Luc*, sh*SMARCA4*, and sh*MAFK* Hi-C data centered on a cluster 3 TE, LTR:ERV1:HERVS71-int on chromosome 19, as the viewpoint showing the flanking 1 Mbp from the center. Both SMARCA4 and MAFK is bound to the TE. Co-binding of SMARCA4 and MAFK is indicated.

(B) As in **panel A**, but centered on an LTR:Gypsy:MamGypLTR3a on chromosome 8.

(C) As in **panel A**, but centered on a SINE:5S-Deu-L2:AmnSINE on chromosome 7.
